# Role of CCK1 receptor in metabolic benefits of intestinal enteropeptidase inhibition in mice

**DOI:** 10.1101/2024.10.17.618852

**Authors:** Kamal Albarazanji, Simon A. Hinke, Cassandre Cavanaugh, Jianying Liu, Stephen Beck, Neetu Shukla, Rong Meng, George Ho, Raul C. Camacho, James Leonard, Andrea R. Nawrocki

## Abstract

Enteropeptidase (EP; enterokinase) is a serine protease that regulates intestinal protein digestion by converting trypsinogen into active trypsin, and thus initiates activation of the pancreatic zymogen cascade. Chronic inhibition of EP and trypsin (EP/T) with camostat (Foipan, FOY-305) or its active metabolite (FOY-251) causes weight loss in obese mice by reducing intestinal protein absorption and suppression of food intake, however, the mechanisms leading to appetite suppression are not well understood. We tested the hypothesis that cholecystokinin (CCK) signaling mediates the anorectic effects of EP/T inhibition using a CCK1R inhibitor (loxiglumide) or CCK1R knockout (KO) mice. Acute treatment with loxiglumide was able to partially reverse FOY-251-induced gallbladder contraction and delayed gastric emptying in mice. Chronic co-administration of loxiglumide reversed FOY-251 mediated effects on food intake and metabolism in diet-induced obese (DIO) mice. Chronic dosing of FOY-251 caused similar reductions in food intake but greater weight loss in CCK1R KO mice compared to wildtype (WT) mice, primarily due to fat mass loss. Pair fed (PF) groups revealed food intake-dependent and -independent mechanisms of weight loss by FOY-251. Notably, FOY-251 treatment induced sustained weight loss, whereas body weight loss rebounded in PF animals. In CCKR1 KO mice, FOY-251 caused greater weight loss, and increased protein calorie loss relative to that in WT mice, while having no effect on glycemic control or FGF21. Hence, CCK1R-dependent and - independent mechanisms modulate the metabolic effects of EP/T inhibition and may play a role in maintaining weight loss by this mechanism.

## Introduction

Obesity is a global epidemic and risk factor for developing type 2 diabetes (T2D) (1). Food intake and energy expenditure are the canonical determinants of body weight. Recent development of SGLT2 inhibitors and other drivers of nutrient uptake underscore that caloric excretion via urine and feces are critical determinants of whole-body energy balance and directly or indirectly interact with central mechanisms of appetite regulation and energy expenditure (2, 3). Enteropeptidase (EP) is an intestinal serine protease responsible for activating the zymogen cascade allowing absorption of amino acids and triglycerides. Congenital EP deficiency is associated with a lean phenotype in humans hence, EP inhibition has been proposed as a potential targeted therapeutic for the treatment of human obesity (4-6). EP is localized on the duodenal brush border, playing a key role in dietary protein digestion by converting inactive trypsinogen into active trypsin (7). Trypsin activates downstream pancreatic zymogens, including chymotrypsinogen, proelastase, and procarboxypeptidases, leading to further protein digestion in the gut (8). We and others have demonstrated that small molecule EP inhibitors cause weight loss and increase fecal protein excretion; furthermore, EP inhibition increases plasma fibroblast growth factor 21 (FGF21) and improves glucose homeostasis in diet-induced obese (DIO) and genetically obese mice (9, 10).

Cholecystokinin (CCK) is a hormone secreted by intestinal I-cells, a subtype of enteroendocrine cells of the proximal small bowel, upon contact with ingested amino acids and fatty acids, with aromatic amino acids and long chain fatty acids being the strongest stimuli (11). Among the major physiological roles attributed to CCK are the regulation of gallbladder contraction, slowing of gastric emptying and inhibition of food intake. CCK evokes these responses by activating either of two cognate G protein-coupled receptors: CCK1R, highly expressed in smooth muscle cells of the gastrointestinal tract, and CCK2R, primarily expressed in the central nervous system (12-16). The actions of peripheral CCK with respect to satiety, gallbladder contraction, and gastric emptying are well documented to be mediated through the CCK1R in humans and rodents (12, 14, 17-19).

Several studies suggest a common role for EP inhibition in food intake and body weight regulation. Camostat, a synthetic EP and trypsin (EP/T) inhibitor, has been shown to reduce food intake and body weight in a variety of obese rodent models (9, 20, 21). Our work identified a novel mechanism of camostat-induced induction of integrated stress response (ISR) genes and FGF21 secretion in DIO and *ob/ob* mice, which may also contribute to metabolic benefits of EP/T inhibition (20). Importantly, it has been shown that camostat stimulates CCK release (21-25), indicating a possible link between EP/T inhibition and the metabolic action of the CCK hormone. However, there is limited information about the role of CCK1R in EP/T inhibition effects on gallbladder and gastric emptying, food intake, and body weight regulation. Our proposed paradigm is that EP/T inhibition drives body weight loss and metabolic improvements through satiety related mechanisms by evoking CCK release from the duodenum, in addition to previously described mechanisms such as stimulation of FGF21 or fecal excretion of protein (9).

The aims of these studies were to: (1) to evaluate the role of CCK1R in the camostat metabolite FOY-251-induced increase in gallbladder emptying, delay in gastric emptying and reduction in food intake in mice using the CCK1R inhibitor, loxiglumide, and (2) to investigate the chronic metabolic effects of EP/T pathway inhibition with FOY-251 in DIO CCK1R knockout (KO) mice.

## Material and Methods

### Animals

Animal studies were performed in compliance with the Animal Welfare Act and protocols were approved by the Institutional Animal Care and Use Committee at Janssen Pharmaceutical R&D (Spring House, PA). Lean 12-week old male C57BL/6 mice were purchased from Taconic Biosciences (Germantown, NY). Obese 16-week old male DIO C57BL/6 mice were purchased from Taconic. Male CCK1R KO mice and their aged-matched wild type (WT) controls on a 129S1/SvImJ background were purchased from the Jackson Laboratory (Bar Harbor, ME). Genotypes were reconfirmed by PCR. WT and KO mice arrived at 8 weeks of age and were housed individually in a temperature-controlled room with 12-hour light/dark cycle with free access to water and a 60% high fat diet (HFD; Diet D12492; Research Diets, New Brunswick, NJ) for 20 weeks to render them DIO.

### Compounds

In plasma, camostat is hydrolysed by carboxyesterase to 4-(4-guanidinobenzoyloxy)phenylacetic acid (FOY-251; camostat metabolite) (26, 27), which has low oral bioavailability (F <2%) and dual potency on Enteropeptidase (EP) and trypsin (T) (k_inact_/K_I_ =9295 and 56340, respectively). For acute studies, FOY-251 was formulated for oral administration in 0.5% hydroxypropyl methylcellulose (HPMC; Sigma-Aldrich, St. Louis, MO), and for chronic studies, FOY-251 was formulated as an admixture in high fat diet (1.59 mg/g), a dose selected based on a previous weight loss study in DIO mice (9). The CCK1R antagonist loxiglumide was purchased from Tocris Biotechne (Minneapolis, MN). Exendin-4 purchased from Bachem (Torrance, CA). Rimonabant (a CB1 receptor inverse agonist), FOY-251 and acetaminophen were supplied by Janssen chemists.

### In Vivo gall bladder contraction assay

Following 20 weeks of HFD feeding with or without FOY-251 formulated in diet (described above), CCK1R KO mice and WT control animals were fasted overnight with free access to drinking water. Fasting body weights were measured and then mice were administered vehicle (5% hydroxypropyl methylcellulose; HPMC; Sigma-Aldrich, St. Louis, MO) or a 100 mg/kg dose of FOY-251 as an HPMC suspension by oral gavage. Mice were euthanized 15 minutes later by CO_2_ asphyxiation. A midline laparotomy was performed, and the gallbladder was removed by using fine forceps to grab the cystic duct to gently remove the gallbladder without disrupting the contents, and gall bladder weight was measured on an analytical balance.

Following a similar protocol, the effect of an oral loxiglumide dose-range on gallbladder emptying was evaluated in overnight fasted lean C57BL/6 mice. Animals were randomized and fasted overnight with access to water. Mice were orally dosed with either vehicle (20% hydroxypropyl-β-cyclodextrin (HP-β-CD) or loxiglumide in the same excipient. Animals were sacrificed 75 minutes post-dose and gall bladder weights measured. In a subsequent experiment, the effect of loxiglumide on FOY-251-induced gall bladder emptying was assessed using the same paradigm, dosing loxiglumide at t=0min, FOY-251 60 minutes later, and collecting gall bladder weights at 75 minutes; a separate group of naïve animals was used to confirm the absence of an effect of the vehicle treatments on gall bladder weights.

### In Vivo acetaminophen gastric emptying assay

To examine the interaction of CCKR1 blockade on the camostat metabolite effect on gastric emptying, the surrogate in vivo acetaminophen appearance assay was tested in DIO C57BL/6 mice. Overnight fasted animals were segregated into groups by body weight. To fully control the experiment, animals were dosed twice separated by 10 minutes: vehicle or 30mg/kg loxiglumide (t=-40min) then vehicle or 100mg/kg FOY-251 (t=-30min). Animals were subsequently injected subcutaneously with phosphate buffered saline (PBS) or 40ug/kg Exendin-4 (t=-20min) and then at t=0min, all animals received an oral dose of 100mg/kg acetaminophen formulated as a suspension in 0.5% HPMC/5% acacia gum (Sigma-Aldrich). Whole blood was collected at intervals indicated for 90 minutes after acetaminophen administration using GE DMPK-C dry blood spot cards (Sigma-Aldrich) and blood acetaminophen quantified by mass spectrometry. Exendin-4, a glucagon-like peptide-1 receptor agonist, was used as a positive control.

### Food intake measurement in DIO mice

The PK parameters of 30mg/kg oral loxiglumide in lean mice (Tmax = 0.5h; Cmax = 8.36uM; t_1/2_ = 3h) were used as the basis for selecting an experimental paradigm measuring food intake over 4 hours to ensure adequate inhibition of CCK1R. DIO mice were maintained on a reverse light cycle and sham dosed for at least 1 week prior to the study. Baseline 4-hour food intake was measured from the initiation of the dark cycle for two sequential days, and the average food intake was used to group animals. On the day of the experiment, body weights were recorded, and animals orally dosed with vehicle or 30mg/kg loxiglumide 30 min before oral dosing the animals with vehicle or 100mg/kg FOY-251, such that the camostat metabolite was given 30 minutes before the animal room lights turned off; food intake over a 4-hour span was measured by hopper weight difference. Rimonabant (3mg/kg) was used as a positive control administered with the same timing as FOY-251, in animals that first received oral vehicle.

### Chronic treatment of CCK1R KO mice with camostat metabolite FOY-251

Male DIO WT and CCK1R KO mice were used. Baseline body composition was measured using Echo MRI (Houston, TX) to randomize mice by body fat mass. Test groups included a control group fed HFD containing no compound, a group fed FOY-251 HFD admixture formulated to deliver approximately 100mg/kg camostat metabolite based upon food intake and body weight from a previous study, and a pair fed (PF) group, provided the same amount of non-medicated food to match that consumed by the FOY-251-treated group. Daily body weight and food intake were measured throughout the 4 weeks of treatment. On day 27, animals were fasted for 5 hours to obtain tail blood glucose (OneTouch Ultra glucometer; Lifescan, Milpitas, CA) and plasma insulin (Rat/Mouse Insulin Assay; MesoScale Discovery, Rockville, MD), followed by body composition analysis using Echo MRI. Cumulative dried fecal samples were collected weekly for fecal protein measurement using the Pierce Detergent Compatible Bradford Protein Assay Kit (Thermo Scientific) as previously described (7). On day 28 of treatment, mice were euthanized by CO_2_ asphyxiation. Blood was collected by cardiac puncture and centrifuged at 10,000rpm for 10 min at 4°C to obtain plasma which was aliquoted into 96-well plates and stored at -80°C until further analysis. Terminal plasma bioanalysis included: FGF21 (rat/mouse FGF21 ELISA Kit, Millipore), amino acids (LCMS, QTrap 5500 Sciex coupled to a Shimadzu prominence LC), lipids and liver enzymes (Alfa Wassermann, West Caldwell, NJ). A partial necropsy was performed to permit liver fat measurement by EchoMRI and pathological assessment of liver histology sections. Neutral buffered formalin-fixed liver sections were stained with hematoxylin and eosin (H&E) using a standardized protocol. Representative digital images were determined by a board-certified pathologist who was blinded to treatment groups. Steatosis was evaluated using semiquantitative grading of liver vacuolation (distinct clear vacuoles of variable size, with displacement of nucleus): minimal (grade 1), mild (grade 2), moderate (grade 3), marked (grade 4) and severe (grade 5). Visiopharm software was used for quantitative image analysis to determine percent vacuolar burden density in total area of vacuoles within the livers.

### Statistical analysis

One-or Two-Way ANOVA followed by Tukey’s or Dunnett’s test for multiple comparisons (with repeated measures for time series data) was used in all studies, as noted in figure legends. An ordinal logistic regression model used for liver vacuolation analysis from histology grading. All tests used the software GraphPad Prism (GraphPad 9). Statistical significance was defined as *p < 0.05.

## Results

### Effect of FOY-251 on gallbladder weights in CCK1R KO mice or after pharmacologic CCK1R blockade with loxiglumide

The average body weight of mice fed high fat diet for 20 weeks prior to use in the gall bladder bioassay were not statistically different (CCK1R KO mice: 48.7±2.4g, n=11 versus WT controls: 49.9±2.31g, n=10). Gallbladders collected from CCK1R KO mice (90±5.9mg) were significantly heavier compared with WT controls (35.5±3.98mg; p<0.05). Treatment with FOY-251 significantly reduced gallbladder weight in both WT (57.7%) and CCK1R KO (19.2%) mice relative to their respective control groups (Figure 1A; p<0.05).

**Figure 1.**
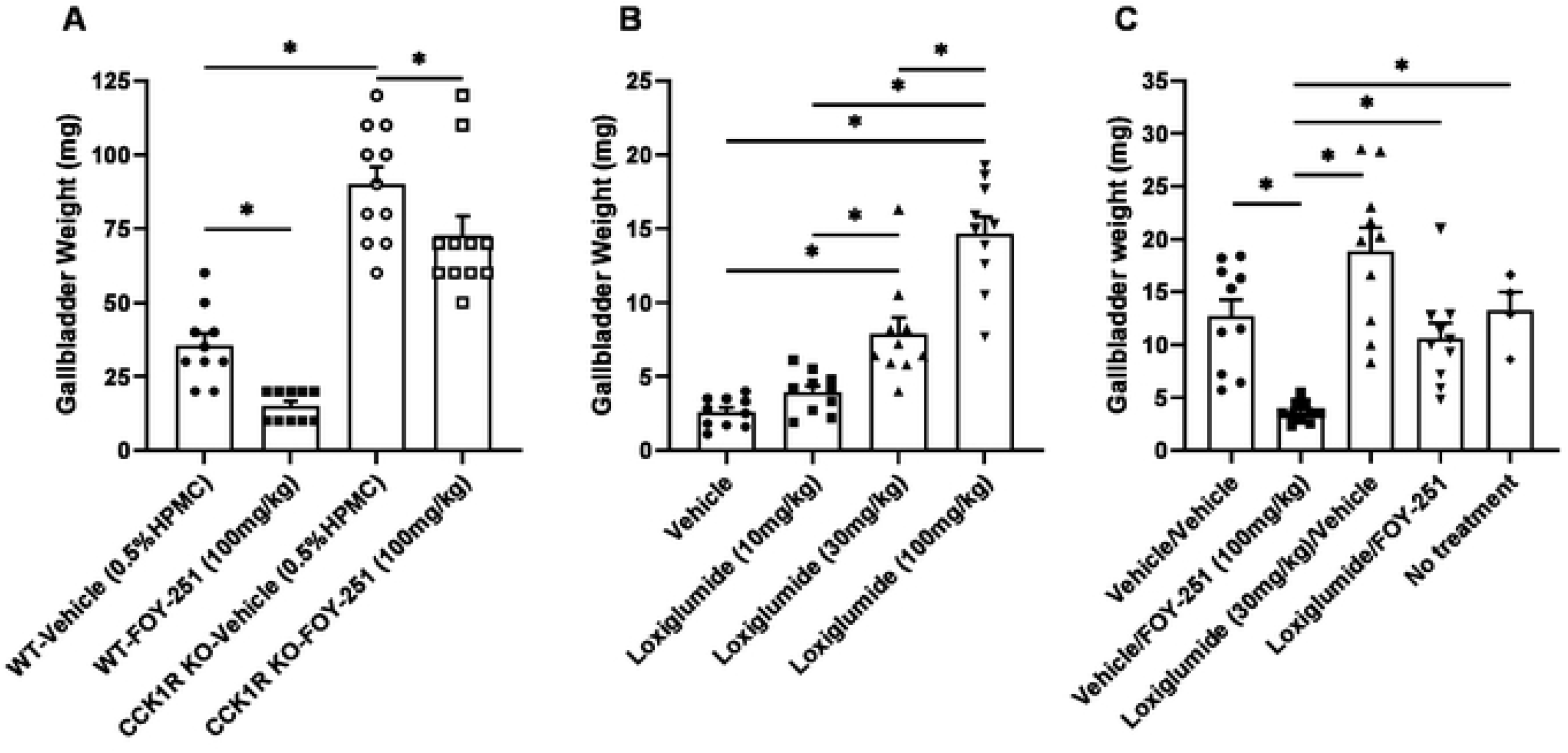
The effect of FOY-251 on gallbladder weights in genetically CCK1R deficient mice or CCK1R blockade with loxiglumide. **(A)** Gallbladder weights 15 minutes after oral vehicle or FOY-251 administration to fasted WT control and CCK1R KO mice. **(B)** Effect of oral loxiglumide at increasing doses on gallbladder weight in fasted C57BL/6 mice. **(C)** Effect of FOY-251 and loxiglumide administration on gallbladder weight in fasted C57BL/6 mice. Animals were first treated with oral vehicle or loxiglumide (t=0min), followed by oral vehicle or FOY-251 (t=30min); gallbladder weight was measured at t=45min. Gallbladders from four fasted naive mice were included to demonstrate an absence of effect of vehicle treatment. Values are expressed as mean ± SEM; *p<0.05 for the indicated comparisons.

Loxiglumide, a potent CCK1R antagonist, dose-dependently increased gallbladder weight in overnight fasted lean C57BL/6 mice (Figure 1B). To examine the effect of CCK1R inhibition on camostat metabolite-induced gall bladder emptying, animals were pre-treated with loxiglumide before administration of oral FOY-251 (Figure 1C). Compared to vehicle treated control animals, FOY-251 reduced gall bladder weight by 71.8% (p<0.01) and loxiglumide increased gall bladder weight by 48.4% (p=0.05). In the group treated with both loxiglumide and FOY-251, gallbladder weight was similar to the vehicle group (10.58±1.4mg vs vehicle 12.70±1.6mg, p>0.05), suggesting that FOY-251 attenuated or reversed the increase in gallbladder weight induced by loxiglumide (Figure 1C). Gallbladder weights in naïve mice were not different than vehicle treated animals.

### CCK1R inhibition attenuates the FOY-251 induced delay in gastric emptying in DIO mice

Protein digestion in the small intestine stimulates CCK secretion which leads to a delay in gastric emptying, induction of gallbladder emptying and inhibition of food intake through CCK1R (9). FOY-251 significantly delayed gastric emptying in DIO mice, evident from reduced blood acetaminophen appearance (AAP; Figure 2A) and AUC_0-90min_ (Figure 2B). Relative to the vehicle-treated control group, FOY-251 reduced gastric emptying (AAP AUC_0-90min_) by 53.4% (p<0.05), whereas CCK1R blockade with loxiglumide caused a non-significant increase in gastric emptying (14.7%; p>0.05) in overnight fasted DIO mice. Pretreatment with loxiglumide partially blocked FOY-251 induced reduction in gastric emptying. The positive control, exendin-4, significantly decreased gastric emptying, consistent with published literature (28) on the effect of GLP-1 receptor agonists on gastric emptying and serves to validate our experimental conditions.

**Figure 2.**
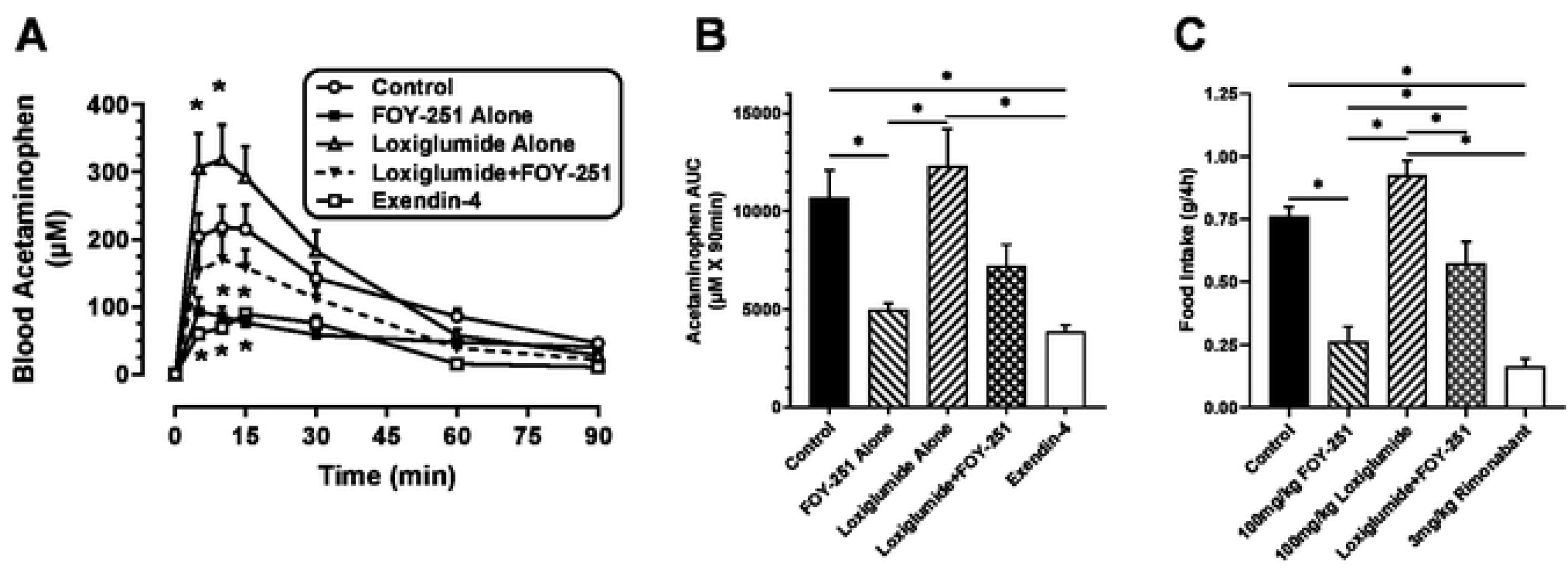
Effect of FOY-251 and/or loxiglumide on gastric emptying and food intake in fasted DIO mice. **(A)** Blood acetaminophen appearance following treatment with 100mg/kg oral FOY-251, 30mg/kg loxiglumide, or their combination (N=7-8/group). 20μg SC exendin-4 was used as a positive control to validate the experimental model (N=S). All animals received oral gavage or subcutaneous injection at the following timepoints: oral vehicle (20%HP-b-CD) or loxiglumide (t=-40min), oral vehicle (5%HPMC) or FOY-251 (t=-30min), subcutaneous PBS or exendin-4 (t=-20min), followed by oral acetaminophen (100mg/kg) at t=0min. *p<0.05 vs vehicle control group. **(B)** Integrated blood acetaminophen for data shown in panel **(A). (C)** 4-hour food intake in DIO mice administered oral vehicle or loxiglumide (t=-60min), followed by oral vehicle, FOY-251 or Rimonabant at t=-30min. Food consumption during the dark phase was measured over 4-hours starting at t=0min (N=8/group). Values are expressed as mean ± SEM; *p<0.05 for the indicated comparisons.

### Loxiglumide impairs FOY-251-induced food intake inhibition in DIO mice

FOY-251 alone significantly decreased 4-hour food intake by 65.6% in DIO mice relative to control animals (Figure 2C). In contrast, CCK1R inhibition with loxiglumide alone trended to increase 4-hour food intake by 21.3% (p>0.05). Pretreatment with loxiglumide limited the FOY-251 induced reduction of food intake to only 24.6% less than control animals (p<0.01). Rimonabant, used as a positive control, significantly decreased 4-hour food intake as expected.

### Chronic effect of FOY-251 on body weight, food intake and fecal protein excretion in CCK1R KO mice

The average daily FOY-251 dose based on food intake over the 4 weeks of treatment was 94.72±2.2 mg/kg and 96.71±1.53mg/kg in CCK1R KO mice and the WT controls, respectively (Figure 3A). Food intake was significantly reduced by FOY-251 treatment on day 1 in both WT and KO mice compared to animals provided non-medicated diet (Figure 3B); daily food intake was lower in FOY-251 diet admixture treated groups through day 12, but was not statistically different thereafter (Figure 3B). Weekly cumulative food intake was significantly reduced by FOY-251 administration during week 1 in both CCK1R KO mice and WT control animals (Figure 3C & 3D). There was a trend for reduced cumulative food intake on week 2, but by weeks 3 and 4 there was no significant difference between FOY-251-treated and control group in either WT or CCK1R KO mice.

**Figure 3.**
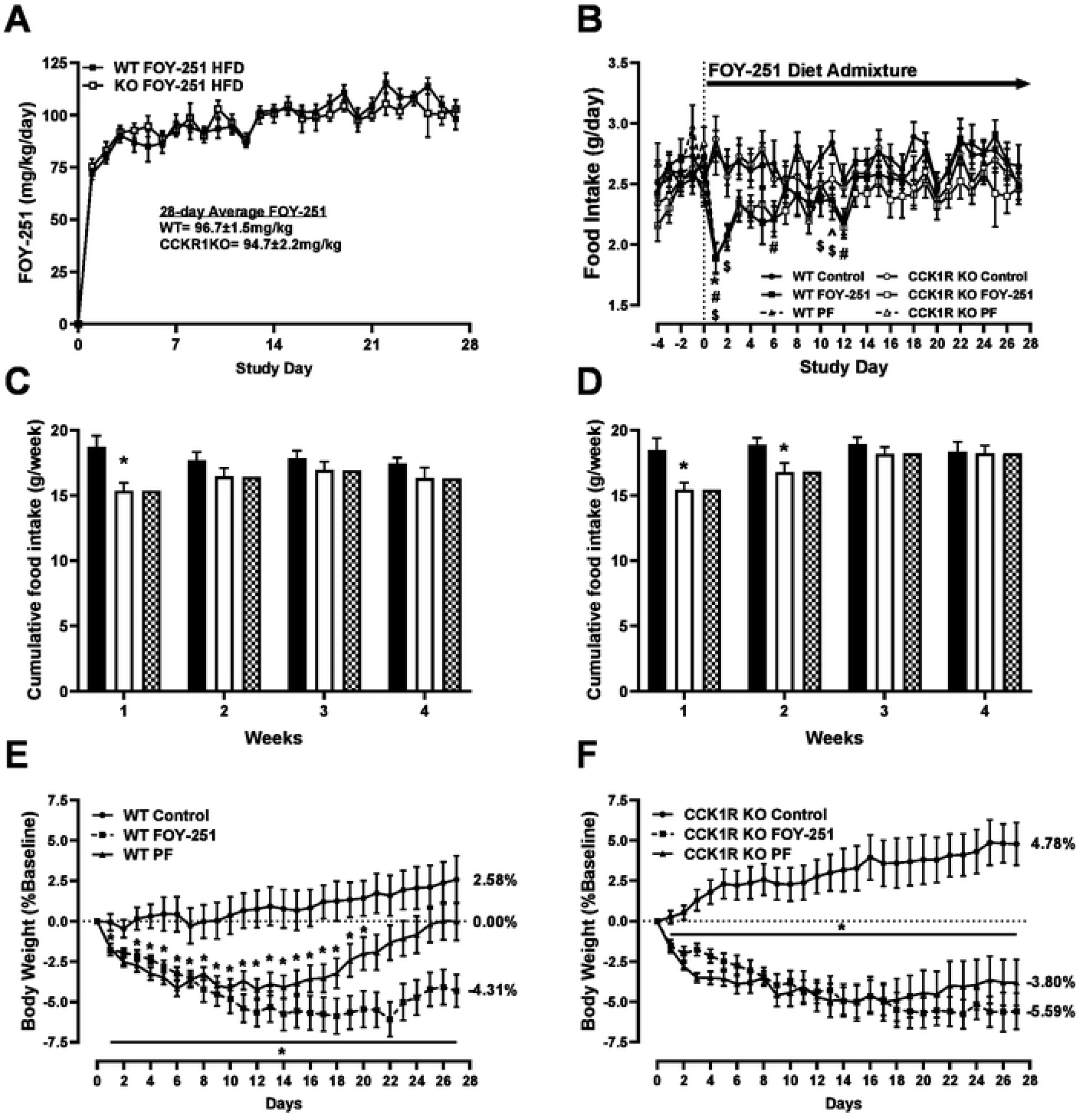
Chronic food intake and body weight effects of FOY-251 administered as high fat diet (HFD) admixture to WTand CCK1R KO. **(A)**Daily FOY-251 compound dose calculated using diet consumption and body weight. FOY-251 was formulated at 1.59mg/gm of 60% HFD. **(B)** Daily food intake WT and CCK1R KO mice provided with non-medicated diet (Control), HFD with FOY-251, and animals pair-fed (PF) non-medicated diet at the same amount as that consumed by respective FOY-251 treated groups. ^^^p<0.05 WT-Control vs WT-FOY-251; ^$^p<0.05 WT-Control vs WT-PF; *p<0.05 CCK1R KO-Control vs CCK1R KO-FOY-251; ^#^p<0.05 CCK1R KO-Control vs CCK1R KO-PF. Weekly cumulative food intake over 4 weeks in wild type mice **(C)** or CCK1R KO mice **(D)** treated with FOY-251 (white bars), or non medicated HFD Control (black bars) or animals pair fed to the same level as FOY-251 treated mice (checkered bars). *p<0.05 Control vs FOY-251 for each genotype. Percent change in body weight in WT **(E)** and CCK1R KO **(F)** mice treated with FOY-251, non-medicated HFD Control or pair-fed (PF) non medicated HFD at the same level as FOY-251-treated group. *p<0.05 vs Control. Data represent Mean±SEM of 9-10 animals per group.

Body weight of WT and CCK1R KO mice fed high fat diet for 20 weeks prior to FOY-251 treatment was 42.6±0.8g (n=28) and 41.1±0.8g (n=29), respectively (p>0.05). Control groups of WT and CCK1R KO mice maintained on non-medicated HFD for the remainder of the study gained 2.6% and 4.8% body weight over baseline (Figures 3E & 3F). Treatment with FOY-251 HFD admixture significantly reduced body weight in both WT and KO mice: a vehicle-adjusted difference of 6.9% (WT) and 10.4% (KO). Pair feeding studies were performed to assess food intake independent effects on weight loss. Notably, WT mice pair-fed the equivalent to the daily food intake of the WT FOY-251 group gradually regained the earlier weight loss (Figure 3E), consistent with the rebound in food intake (Figure 3B and 3C). In contrast, the pair-fed CCK1R KO mice did not show a similar return of body weight towards baseline that was observed in pair fed WT animals (Figure 3F).

Body composition is presented as difference between the baseline and day 27 of treatment for each individual animal (Figures 4A & 4B). Fat mass and lean mass were not different comparing control non-medicated HFD groups of WT and CCK1R KO mice. Chronic FOY-251 treatment reduced fat mass by 14.4% in WT mice (p<0.01) and by 21.2% in KO animals (p<0.0001) (Figure 4A). Consistent with body weight data presented in Figure 3, pair-fed WT animals showed no difference in fat mass relative to control WT animals after the 4-week study period, whereas pair-fed CCK1R KO mice had significantly lower fat mass than controls of the same genotype (Figure 4A). FOY-251 had no effect on lean mass in either CCK1R KO mice or WT controls relative to non-medicated diet fed control groups. However, pair-fed groups of both genotypes showed a small reduction in lean mass (Figure 4B).

**Figure 4.**
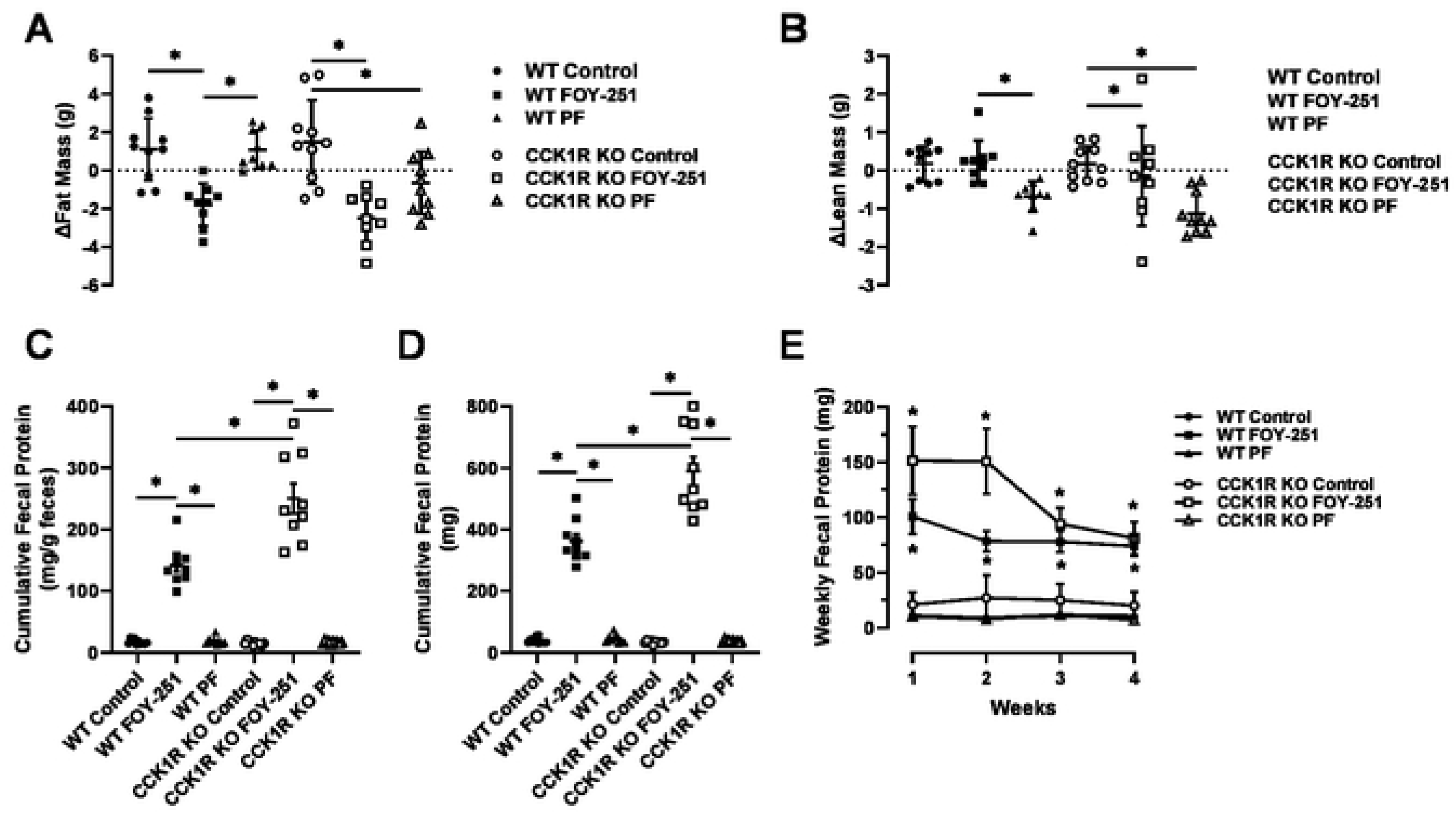
Effect of chronic FOY-251 treatment on body composition and fecal protein excretion in WT and CCK1R KO diet-induced obese mice. Body composition of WT and CCK1R KO mice fed control HFD, FOY-251 HFD admixture, and animals pair-fed to the FOY-251 group was measured by whole body MRI to measure fat mass **(A)** and lean mass **(B)**. Data represent change in body composition from baseline for each individual animal. Cumulative protein in excreted feces over the 4 weeks of treatment was normalized to total fecal output **(C)** or presented as absolute total protein excreted **(D)** in WT and CCK1R KO mice receiving non-medicated high fat diet, FOY-251 HFD admixture or pair fed non-medicated HFD. **(E)** Weekly cumulative total fecal protein excretion over the 4-week study period in CCK1R KO mice and WT controls. Data represent the mean±SEM of 9-10 animals per group; *p<0.05 for the indicated comparisons.

FOY-251 inhibition is associated with enhanced fecal protein excretion in obese mice (9). Unlike food intake, treatment with FOY-251 maintained elevated fecal protein excretion (caloric loss) throughout all 4 weeks of treatment in both CCK1R KO mice and WT controls (Figures 4C, 4D, and 4E). There was no difference in fecal protein excretion expressed either as fecal protein concentration (mg/g feces) or total fecal protein content (mg) between control animals of each genotype, or between pair-fed groups and their respective controls. Treatment with FOY-251 significantly increased both cumulative fecal protein concentration and total protein content in both WT and CCK1R KO animals. WT FOY-251 treated mice showed 8- to 9-fold greater protein excretion, whereas FOY-251 treatment of CCK1R KO rodents was observed to produce a greater effect (∼16.5-fold increase in cumulative fecal protein concentration and 17.5-fold increase in total fecal protein content; Figures 4C and 4D). Weekly fecal protein content (Figure 4E) was higher in CCK1R KO mice compared with their WT controls. Both CCK1R KO mice and WT controls treated with FOY-251 had significantly higher weekly fecal protein content compared to their respective controls or pair-fed groups (Figure 4E). The lack of an increase in fecal protein in the pair-fed groups supports the conclusion that the increase in fecal protein was FOY-251 mediated.

Terminal liver fat content was assessed by EchoMRI in WT and CCK1R KO mice, showing no difference due to genotype in non-medicated diet fed control groups (Figure 5). Relative to respective vehicle controls, FOY-251 treatment significantly reduced liver fat mass in both genotypes by similar magnitudes (Figure 5A). Liver fat mass in pair-fed CCK1R KO mice were significantly reduced compared with CCK1R KO mice controls (p<0.05), however, a similar reduction was not observed in the pair-fed WT animals. Histopathology scoring the severity of steatosis is presented in Figure 5D, which showed results that were consistent with Visiopharm quantitative image analysis calculating vacuolar area (Figure 5C). There was no significant difference between WT and CCK1R KO mice comparing either histopathology scoring using ordinal logistic regression model analysis or statistical comparison of quantitative image analysis values. FOY-251 significantly decreased the severity of vacuolation in both genotypes of mice (Figures 5C and 5D). A trend towards improvement in steatosis was also observed in pair-fed animals, but to a lesser degree than FOY-251 treated mice (p>0.05 compared to non-medicated controls).

**Figure 5.**
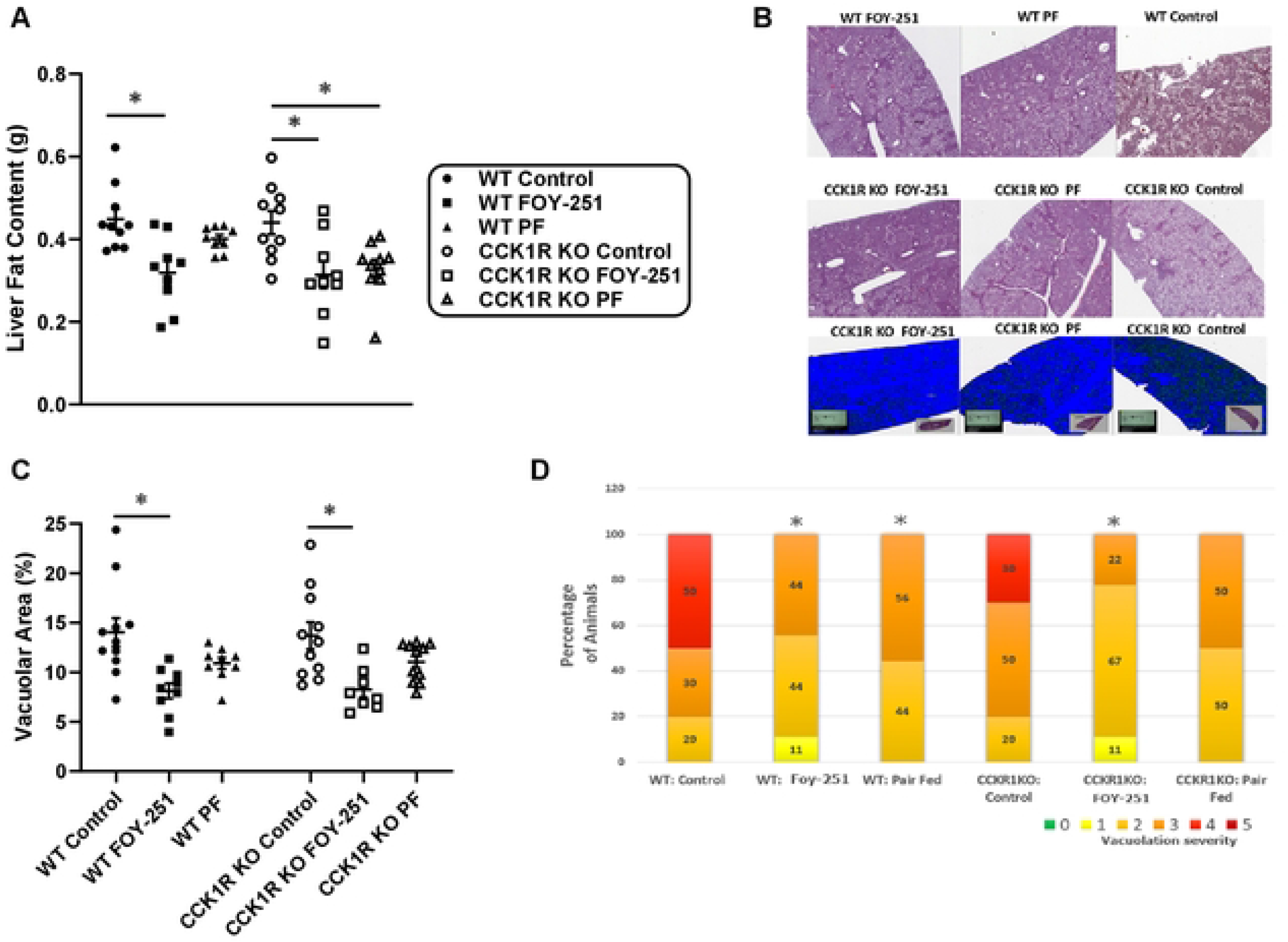
Liver steatosis in chronic FOY-251-treated DIO CCK1R KO mice and their wild type controls. **(A)** Post-mortem organ fat content measured using MRI in WT and KO mice fed control HFD, FOY-251 HFD admixture, and animals pair-fed to the FOY-251 group. **(B)** Representative hematoxylin & eosin stained liver tissue sections from WT and CCK1R KO mice; blue sections indicate the software analysis algorithm determined tissue area for representative CCK1R KO samples. **(C)** Quantitative image analysis for steatosis measuring vacuolar burden (%vacuole area/liver area) using Visiopharm software. **(D)** Semiquantitative blinded histopathology grading of liver vacuolation (distinct clear vacuoles of variable size, with displacement of nucleus) consistent with lipidosis (steatosis) severity: [minimal (grade 1), mild (grade 2), moderate (grade 3), marked (grade 4) and severe (grade 5)). *p<0.05 for indicated comparisons.

No difference in fasting blood glucose or plasma insulin was observed comparing either WT or KO animals, or with FOY-251 treatment (Figures 6A and 6B). In contrast, animals pair-fed to the same amount of HFD that respective FOY-251 animals consumed, showed significant improvement in fasted glycemia (p<0.01) but no change in insulinemia for either genotype of mice. Terminal fed plasma FGF21 was not different between CCK1R KO mice (13.46.7±191.8pg/mL) and their WT controls (1764.5±250.6pg/mL; p>0.05), nor did treatment with FOY-251 cause an effect on FGF21 (Figure 6C). Terminal fed plasma analysis for a panel of amino acids revealed no significant changes in any of amino acids comparing WT to CCK1R KO, with or without FOY-251 treatment, or in pair-fed animals (Supplementary Table 1). No statistical differences were observed by genotype or treatment on terminal plasma liver enzymes (ALT and AST), β-hydroxybutyrate (BHB), or lipid profile (Supplementary Table 2).

**Figure 6.**
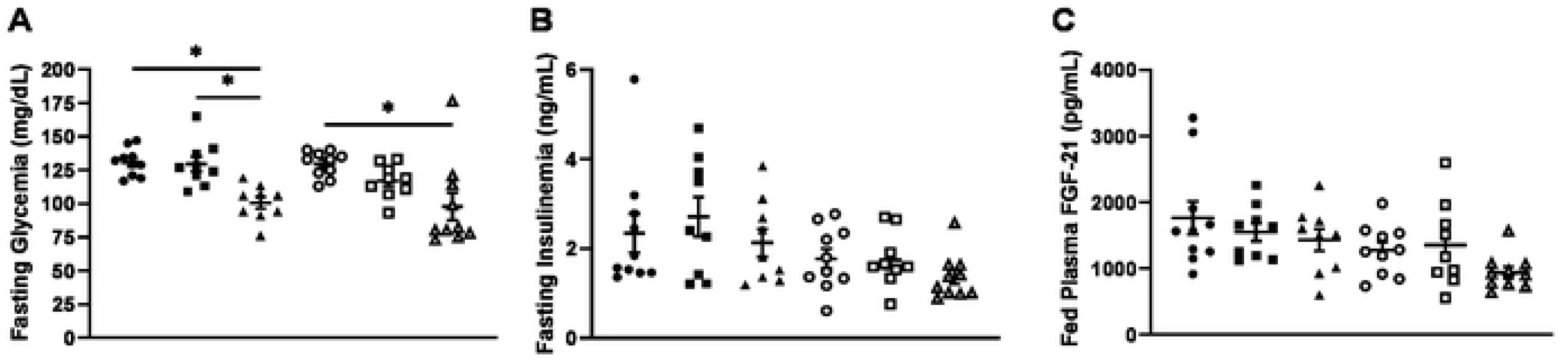
Glucose homeostasis in chronic FOY-251-treated DIO CCK1R KO mice and their wild type controls. Fasted blood glucose **(A)** and plasma insulin levels **(B)** were obtained on day 27 of treatment following a 5-hour fast in WT mice (filled symbols) and CCK1R KO mice (open symbols) given non-medicated high fat diet (circles), HFD formulated with FOY-251 (squares), or animals pair-fed non-medicated diet (triangles) at the same amount consumed by FOY-251 treated mice. *p<0.05 for indicated comparisons. **(C)** Plasma FGF-21 was measured in terminal fed plasma samples taken during sacrifice after 4-weeks of treatment. Data represent mean±SEM of 9-10 animals per group.

## Discussion

Intestinal protein digestion leads to pancreatic zymogen secretion that drives CCK release, which modifies gastric emptying, gall bladder contraction and promotes satiety. In this study, we examined the interplay between digestive serine protease activity and the CCK1R pathway in regulating gastric motility, gall bladder contraction, food intake and associated metabolic benefits. We describe the physiological response of inhibiting EP/T in obese mice in presence and absence of a functional CCK1R response using genetic (CCK1R KO mice) using a pharmacological tool (FOY-251, the active metabolite of camostat). Our data demonstrate that EP/T inhibition leads to an accumulation of fecal protein and excretion in DIO mice and has profound effects on gallbladder weight, gastric emptying, and food intake, leading to weight loss in DIO mice through CCK1R signaling. We show that the chronic EP/T inhibition by FOY-251, drives acute anorectic effects, long term weight loss and metabolic improvements through CCK1R pathway related mechanisms. In contrast, it appears that acute FOY-251 effects on gallbladder and gastric emptying are only partially reversed in the absence of functional CCK1R, indicating CCK-independent mechanisms. CCK is a major controller of gallbladder contraction and has been shown to decrease fasting gallbladder volume in humans and mice (29, 30). In our study, FOY-251 significantly decreased gallbladder weight in both CCK1R KO mice and WT controls (Figure 1A), suggesting that FOY-251 can mediate gallbladder emptying independently of CCK1R. This was also evident from the gallbladder weights of CCK1R KO mice which were significantly heavier than those from WT controls (Figure 1A). This is consistent with a previous report showing exogenous CCK administration caused bile acid secretion in bile duct cannulated CCK1R (+/-) and CCK1R (+/+) mice but not in CCK1R KO mice (19). However, the partial reversal of gallbladder emptying in overnight fasted C57BL/6 mice induced by FOY-251 in the presence of loxiglumide, may suggest that in addition to its direct effect on gallbladder emptying, EP/T inhibition through the stimulation of endogenous CCK release also influences gallbladder emptying. The dose-responsive increase in gallbladder weight by loxiglumide also suggests that there is a fasting CCK1R tone without CCK stimulation upon ingestion of nutrients, or as in this study stimulation of CCK release through administration of FOY-251 in C57BL/6 mice (Figure 1B). In fasted subjects, blockade of CCK1R with loxiglumide increased gallbladder volume, supporting that in addition to inducing postprandial gallbladder contraction, CCK is also involved in fasting gallbladder volume regulation (30).

Another physiological role of CCK is to slow gastric emptying. Infusion of CCK to levels similar to physiological postprandial concentrations inhibited gastric emptying of liquid and semisolid meals in normal subjects, and blocking CCK1R with loxiglumide significantly accelerated gastric emptying of both liquid mixed meals and glucose (17). In rats with implanted gastric fistulae, inhibition of CCK1R with synthetic CCK1R antagonist L364,718 blocked the delay in gastric emptying induced by exogenous infusion of CCK-8 or that induced by endogenous CCK secretion stimulated with peptone solution or camostat (31). In our study, FOY-251 significantly delayed gastric emptying in fasted C57BL/6 mice, while loxiglumide accelerated gastric emptying (Figures 2A and 2B). Blocking CCK1R with loxiglumide partially reversed the delay in gastric emptying induced by FOY-251 (Figures 2A and 2B), suggesting that EP/T inhibition evokes endogenous CCK secretion, and results in reduced gastric emptying via CCK1R. Since loxiglumide alone increased gastric emptying in fasted mice, this confirms the contribution of endogenous CCK in the regulation of gastric emptying.

With respect to the role of CCK1R in acute food intake, we found that FOY-251 caused a significant reduction in 4-hour food intake in DIO mice, and this effect was fully reversed by blocking CCK1R with loxiglumide (Figure 2C). This is consistent with data showing acute reduction of food intake in rats by camostat and reversal by CCK1R antagonist devazepide, but not by CCK2R antagonist L365,260 (32). Measuring Fos-like immunoreactivity in fasted rats, it was demonstrated that the effect of camostat on acute food intake is likely mediated by the vagus nerve, specifically by activation of the dorsal vagal complex but not the myenteric plexus (32).

Chronic FOY-251 treatment in WT and CCK1R KO mice revealed an interesting role of CCK1R signaling associated with the activity of EP/T enzymes in the control of fecal protein excretion, food consumption and body weight regulation. We observed no difference in food intake comparing WT mice to CCK1R KO animals fed a high fat diet (Figures 3B-D), consistent with a previous report showing no difference in food intake between HFD-fed CCK deficient mice and WT controls (33). When comparing the effect of chronic FOY-251 treatment of obese WT and CCK1R KO mice, our data suggest the existence of food-intake independent mechanisms involved in weight loss, and that CCK1R deficiency plays a role in the maintenance of weight loss induced by inhibition of EP/T. FOY-251 administration caused a rapid inhibition of food intake which waned after approximately two weeks in both WT and CCK1R KO mice. FOY-251 caused sustained weight loss over the 4-week treatment period whereas pair-fed counterpart animals experienced weight regain during the same time implicating mechanisms beyond adaptation to caloric restriction associated with this mechanism. This rebound effect of body weight with pair-feeding was not observed in animals lacking CCK1R (Figures 3E and 3F) suggesting that CCK release and its effect on food intake wane after chronic weight loss in this model. SCO-792, a synthetic EP inhibitor, induced similar reduction in food intake during only the first 2 weeks of treatment in DIO mice (10). Although CCK agonism alone is ineffective at eliciting or maintaining weight loss, when co-administered with amylin or amylin plus leptin, it augmented the durability of food intake and body weight-lowering effects in DIO rats (34). Similarly, co-agonism of GLP-1R and CCK1R using a novel fusion protein consisting of sequence based upon exendin-4 linked to a CCKR1-selective peptide was synergistic, producing ∼28% weight loss over 10 days in DIO mice (35). However, CCK1R agonists have not been effective thus far in human trials as a monotherapy (36, 37). Our data in DIO mice suggest that a CCK1R antagonism in combination with EP/T inhibitors may have therapeutic potential for sustained body weight loss, but translation of this effect to humans remains to be demonstrated.

We and others have provided evidence that the caloric loss through fecal protein excretion induced by synthetic EP/T inhibitors plays a significant role in body weight regulation in DIO mice and *ob/ob* mice (9, 10). Jia et al (38) reported that chronic treatment with camostat increased fecal protein excretion and reduced body weight without effect on food intake in Otsuka Long-Evans Tokushima Fatty (OLETF) rats, a type 2 diabetic rodent model lacking CCK1R expression. To the best of our knowledge, data presented here is the first to evaluate the chronic effect of EP/T inhibition by FOY-251 on fecal protein and weight regulation in CCK1R KO mice and WT controls. Although fecal protein excretion was similar between CCK1R KO mice and their WT control mice, FOY-251 treatment caused a greater fecal protein (calorie) loss in the CCK1R KO mice compared with WT controls (Figures 4C-E) fed on a HFD; this effect was independent of food intake, suggesting a possible role of CCK1R on protein digestion and/or absorption in the gut. The increase in fecal protein (calorie) loss throughout the 4-week treatment with FOY-251, may suggest that fecal calorie loss contributes to the body weight loss and the absence of weight regain despite food intake returning to pre-treatment levels after 2 weeks in both CCK1R KO mice and WT controls. Recently, it was suggested that fecal caloric loss is an important contributor to energy balance and weight regulation in humans (2).

In this study, FOY-251 caused no improvement in fasted insulin or glucose and did not increase fed plasma FGF-21 levels in WT or CCK1R KO mice (Figure 6). This is not in agreement with the improvement in plasma insulin/glucose or increase in plasma FGF-21 levels reported by us and others in DIO mice or ob/ob mice (9, 10). The reason for this discrepancy is not clear, despite FOY-251 treatment causing similar effects on reducing body weight and increased fecal protein excretion as observed previously (9). One likely possible explanation is that the mouse strain background used in the current study was different (129S1/SvlmJ) than the C57BL/6 mice used in the earlier report (9). Bi et al (39) reported plasma glucose in DIO WT and CCK1R KO mice on 129/SvEv background; they found that the fed glucose levels were not statistically different between KO mice (172±10.2 mg/dL) and WT controls (198±13.8mg/dL). Similarly, although somewhat lower in magnitude, 5-hour fasted blood glucose levels were not different between genotypes (WT: 130.8±3.21; KO: 129.6±3.02mg/dL; Figure 6A). Burgess and colleagues (40) reported that total body glucose production was ∼30% less in 129S1/SvImJ mice relative to ICR, FVB/N, C57BL/6J mice strains during prolonged fasting. Another example supporting metabolic differences among mouse strains was observed during glucose tolerance testing animals treated with tamoxifen, showing improved glucose tolerance in BALB/cJ and C57BL/6J mice but not in male 129S1/SvImJ mice, suggesting that the strain background can affect glucose homeostasis (41).

Rodent non-alcoholic steatohepatitis (NASH) models can be developed by overnutrition using modified diets (e.g. Western diet, Gubra-Amylin NASH diet, et cetera); the continuum between steatosis and the lipotoxic effects causing heptatocellular damage and lobular inflammation vary depending on the genetic background of the rodent and the specific components of the NASH-inducing diets (42). The potential role of EP/T inhibition and the role of CCK1R in HFD-induced steatosis in 129/SvEv mice was assessed by liver MRI, histopathology scoring and digital image analysis quantifying vacuolation (Figure 5). Liver fat content and percent vacuolar burden (total area of vacuoles within the liver) was significantly reduced by FOY-251 in both CCK1R mice and WT controls. We previously reported significant reduction in liver weight and lipidosis grade in *ob/ob* mice treated with camostat for 7 days (9), and others showed that chronic EP inhibition with SCO-792 significantly reduced hepatic triglyceride and total cholesterol in DIO mice (10). Pair-fed groups of animals receiving the same amount of HFD as that consumed by FOY-251 admixture groups showed smaller trends in liver fat and vacuolation (p>0.05), and were not different in animals lacking CCK1R, suggesting that the effect of FOY-251 on liver steatosis is independent of CCK1R signaling.

In conclusion, EP/T inhibition mediates its biological effects through CCK1R-dependent and - independent mechanisms on gallbladder and gastric emptying. CCK1R plays a key role in acute but not in chronic reduction of food intake induced by EP/T inhibition in DIO mice. The CCK pathway appears to play a role in the body weight rheostat, regulating food intake independent weight gain. Finally, CCK1R appears to play an important role in fecal protein (caloric) excretion induced by EP/T inhibition which plays a role in EP/T induced weight loss. Hence, EP/T inhibitor based therapeutic approaches may be used in combination with approved anorectic agents due to non-redundant mechanisms of action.

## Notes

CONFLICT OF INTEREST Authors are/were employees of Janssen Pharmaceutical Companies of Johnson & Johnson and receive(d) salaries and stock commensurate with employment at the time this research was conducted.

FUNDING This study was supported by Janssen Research & Development.

### Competing Interest Statement

I have read the journal's policy and the authors of this manuscript have the following competing interests: all authors are or were employees of Janssen Pharmaceutical Companies of J&J all authors hold J&J stock.

